# A Comparison of Computational Methods for Modeling Stochastic Collaborative DNA Methylation Dynamics

**DOI:** 10.1101/2025.11.25.690568

**Authors:** Maximillian A.G. Lemus, Elizabeth L. Read

## Abstract

DNA methylation is a widespread epigenetic modification that is important in gene regulation, cancer, and aging. Methylation occurs primarily on symmetric cytosine-phosphate-guanine (CpG) dinucleotides in vertebrates. Recent findings support that the enzymatic reactions governing both addition and removal of DNA methylation at an individual CpG site are influenced by methylation levels at other CpG sites in the vicinity. Mathematical models that treat this phenomenon have been termed “collaborative” models. These models are computationally challenging due to rare switching events and the combinatorial explosion of site-to-site interactions, hindering integration with data. We compare the efficiency and accuracy of three collaborative DNA methylation modeling approaches: a stochastic simulation algorithm (SSA), an exact Chemical Master Equation (CME), and a mean-field CME. The exact CME model is limited to very small system sizes (few CpGs), with computational expense scaling as ∼ 3^2^*^N^* for a system of *N* sites. The SSA approach can accurately handle larger system sizes, with computational expense scaling as ∼ *N* ^2^. The mean-field CME model accurately captures qualitative methylation patterns and is efficient for small (N<100) system sizes, but accuracy and efficiency suffer at larger system sizes, with expense scaling as ∼ *N* ^5.5^. Using the developed numerical approaches, we compute methylation phase diagrams, which reveal how methylation levels and bimodality depend on the DNA-sequence-architecture of CpG clusters, as well as on enzymatic model parameters. We also present phase diagrams derived from the mouse epigenome. These data reveal a complex dependence of cluster methylation patterns on cluster architecture, which is nevertheless well-captured by the relatively simplified model.

## 1. Introduction

DNA methylation is a ubiquitous epigenetic mark that plays important roles in gene regulation, including in the establishment and maintenance of cellular identity [1]. In the mammalian genome, DNA methylation occurs primarily at the 5th carbon of cytosine in cytosine-phosphate-guanine (CpG) dinucleotides, resulting in 5-methylcytosine (5mC). While approximately 80% of CpGs are methylated, methylation patterns vary across genomic regions, across cell types, and across time. In certain areas, CpGs form denser-than-usual clusters called CpG islands that tend to be unmethylated; however, methylated islands near promoters are associated with gene silencing [2, 3]. Changes in DNA methylation patterns are a biomarker of age [4], and aberrant DNA methylation patterns are associated with various diseases, including cancer [5, 6].

The methylation and demethylation of CpG sites, catalyzed by various enzymes, underlies the core dynamics of the DNA methylation system (reviewed in [3, 7]). Methylation is catalyzed by DNA methyltransferases (DNMTs)–the so-called “writers”–while loss of methylation can occur either actively (catalyzed by a group of “eraser” enzymes called ten-eleven translocation (TET) enzymes [8]) or passively, due to DNA replication. Passive demethylation due to replication is normally counteracted by the “maintenance” DNMT1, which has high catalytic activity at hemimethylated sites [9]. Other methyltransferases, namely DNMT3a and DNMT3b, are known as the *de novo* writers, which can establish new methylation at previously unmodified sites. More recent evidence indicates that the roles of the DNMTs are not completely distinct [10].

A relatively simple model of these enzymatic processes emerged in the late 20th century [11, 12, 13]. In this model, (now referred to as the “classical” [10] or “standard” [14] model), each CpG is treated as being in one of three states: unmethylated on both strands (‘*u*’), hemi-methylated (‘*h*’, denoting a methylated 5mC on only one side of the symmetric CpG dinucleotide), or fully methylated (‘*m*’). Each CpG transitions independently between states as a result of the enzymatic reactions, in addition to replication-associated passive loss of methylation. However, this model cannot fully account for features of real mammalian methylation landscapes [10, 15, 16]: it does not explain spatial variation in methylation patterns (most CpGs tend to be methylated, but dense CpG islands are “protected” from *de novo* methylation), and it does not explain how these methylation patterns remain stable despite stochastic variability. A feature of several more recent mathematical models is the interaction, or “collaboration” [14], of CpGs, such that the state of one CpG can influence other CpGs in its local vicinity, through a variety of coupling mechanisms, such as enzyme processivity [9, 17, 18, 19]. These interactions appear necessary to stable methylation patterning [20, 14].

There is recent increased interest in utilizing mathematical models to analyze large epigenomic datasets such as those obtained by Whole Genome Bisulfite Sequencing [21, 19]. However, there are several computational challenges associated with computing methylation distributions from collaborative models, and thus by extension, with parameterizing such mathematical models based on big data. First, the interactions in a collaborative model necessitates treatment of dynamics over a group of CpGs. Even the relatively simplified 3-state (*u, h, m*) description leads to a combinatorial explosion of distinct possible states over a CpG cluster of size *N*. Second, the inherently stochastic nature of individual biochemical reactions necessitates a probabilistic model over discrete biochemical states. The collaborative DNA methylation model is amenable to established approaches from stochastic chemical kinetics, such as the Stochastic Simulation Algorithm (SSA) [22] or the Chemical Master Equation (CME) approach [23, 24]. However, both of these general approaches have drawbacks, namely, long simulation times for SSA, and impractically large system sizes for CME.

In this work, we present a detailed study of the efficiency and accuracy of alternative approaches to modeling stochastic dynamics of mammalian DNA methylation. We focus on a simplified version of a stochastic model originally developed by Haerter et al. [14], which was shown to recapitulate experimental observations on bimodal methylation patterning, including the dependence of methylation levels on the local CpG topology [25]. Based on this general collaborative framework, we recently developed an efficient, mean-field CME approximation [26], which quantitatively recapitulated the switch-like dependence of methylation bimodality on CpG density and the shifts in methylation distributions that result when key enzymes are knocked out. While this mean-field approximation is more amenable to parameterization based on data, it is incapable of predicting methylation at individual CpG sites, and the accuracy of the approximation has not been studied. Therefore, in this study we explore the tradeoffs between efficiency and accuracy of the exact stochastic biochemical model (for which dynamics can be simulated) versus the approximate mean-field model (for which dynamics can be efficiently computed).

Although some version of the collaborative model has been applied in a number of studies [25, 17, 27, 28, 18, 19, 26], the stochastic model’s behavior over the parameter space has not been comprehensively explored. Therefore, in addition to exploring the efficiency and accuracy tradeoff, here we also present phase diagrams that reveal how bimodality and methylation levels depend on enzymatic parameters and sequence-encoded CpG positions, in comparison to real-world methylomes.

## 2. Methods

### 2.1. Overview of the collaborative DNA methylation model

The model is comprised of a 1-dimensional lattice of *N* CpGs with inter-CpG distances *d*. Each site can transition stochastically between *u*, *h*, and *m* states as shown in the schematic, Fig. 1A. Standard methylation reactions comprise the transitions from *u* → *h* and *h* → *m* that occur on a site, independently of other sites. For simplicity, these processes are assume to occur at the same rate, denoted *St*^+^. (Or, more precisely, at the same constant stochastic propensity, defined as the probability of the reaction event occurring per unit time [22, 23]). Likewise, standard demethylation reactions, occuring at rate *St*^−^, comprise independent *m* → *h* and *h* → *u* transitions. (See also Table 1). Collaborative reactions occur at a rate that depends on the current state of surrounding CpGs. In our model, collaboration leads to positive feedback, i.e., nearby CpGs in the *u* state promote demethylating reactions, while nearby CpGs in the *m* state promote methylating reactions. Thus, *Co*^+^ denotes reactions from *u* → *h* and *h* → *m*, influenced by a nearby fully methylated CpG in *m*. Similarly, collaborative demethylation rates are denoted *Co*^−^ and describe reactions from *m* → *h* and *h* → *u*, influenced by an unmethylated CpG in *u*.

**Figure 1:**
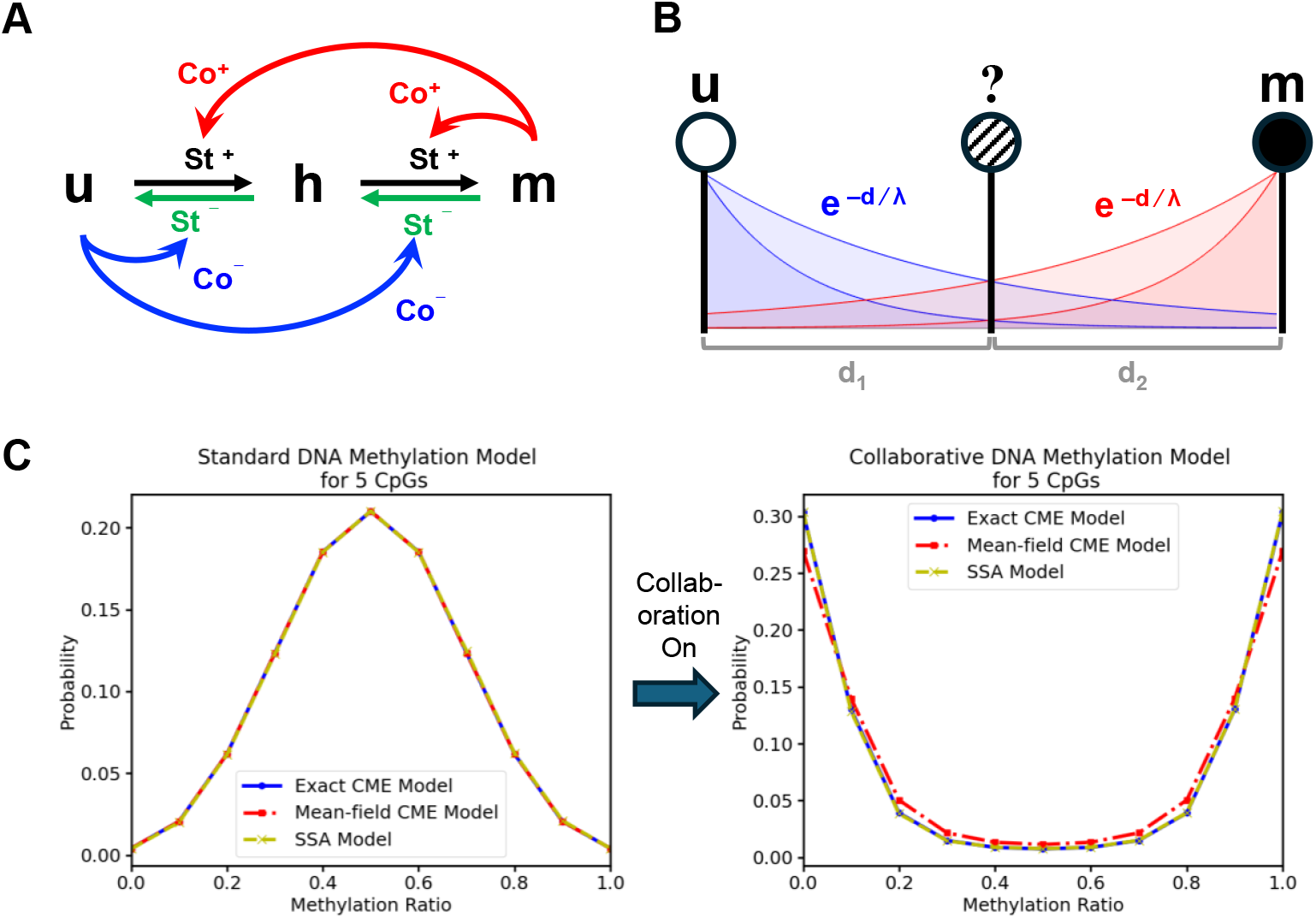
**(A)** Collaborative DNA methylation mechanism with standard methylation reactions *St*^+^, collaborative methylation reactions *Co*^+^, standard demethylation reactions *St*^−^, and collaborative demethylation reactions *Co*^−^. All rate parameters are in units of inverse time *τ* ^−1^ (arbitrary time units, see Methods). **(B)** The exponential distance-dependent field effects for each *u* and *m* site on a neighboring site distances *d*_1_ and *d*_2_ basepairs (bp) away with unmethylated and fully methylated length-scale parameters *λ_u_*and *λ_m_* (units bp^−1^), respectively. **(C)** Probability versus methylation ratio between exact CME, mean-field CME, and SSA models for 5 CpG sites leads to bistability after turning on collaborative DNA methylation rates. For both left and right plots the distance parameter was *d* = 10 and the standard reactions were set to *St*^+^ = *St*^−^ = 1. For the left plot, collaborative reactions were set to *Co*^+^ = *Co*^−^ = 0 and for the right plot, collaborative reactions were set to *Co*^+^ = *Co*^−^ = 5.

**Table 1:**
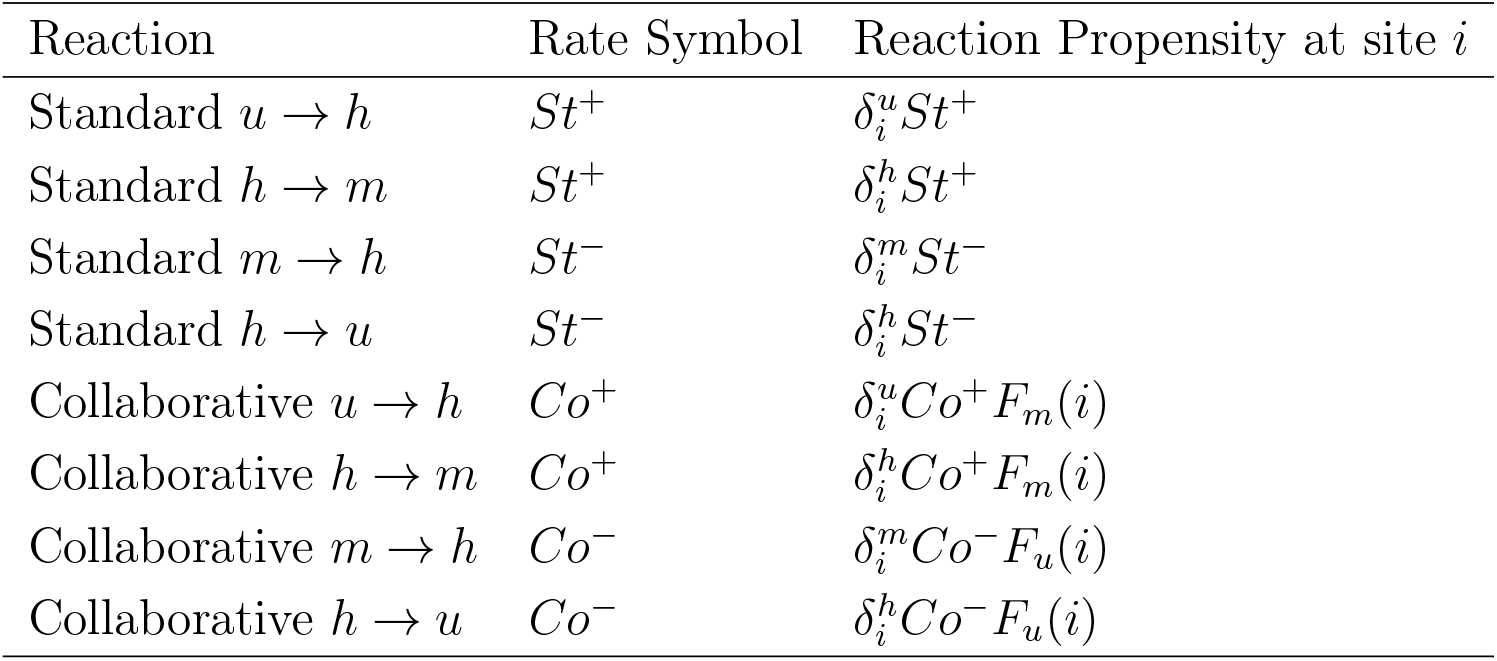
Summary of reaction rate constants and stochastic propensities for the collaborative DNA methylation model. To simplify the parameter space, we assume that standard (de-)methylating reactions occur at the same rate, (*St*^−^) *St*^+^. Similarly, we assume that collaborative (de-)methylating reactions occur at the same rate, (*Co*^−^) *Co*^+^. The propensity for a reaction to occur at a site *i* depends on the site being in the appropriate state, e.g., site *i* must be in *u* state for non-zero propensity of the *u* → *h* reaction, denoted by a delta function δ*^u^_i_*. Collaborative propensities also depend on the methylating (*F_m_*(*i*)) or demethylating (*F_u_*(*i*)) fields exerted by neighbors on site *i*.

**Table 2:**
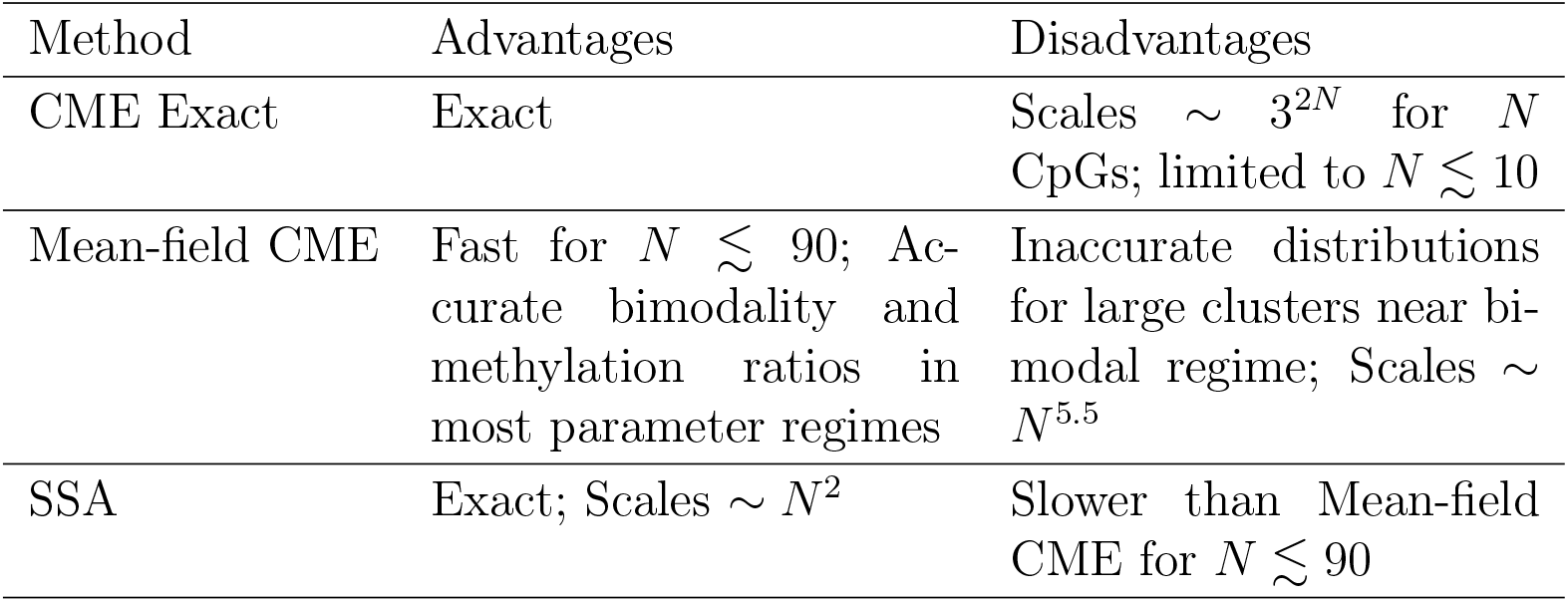
Summarizing differences between all three stochastic computational approaches to the collaborative DNA methylation models.

| Method | Advantages | Disadvantages |
| --- | --- | --- |
| CME Exact | Exact | Scales $\sim 3^{2N}$ for $N$ CpGs; limited to $N \lesssim 10$ |
| Mean-field CME | Fast for $N \lesssim 90$; Accurate bimodality and methylation ratios in most parameter regimes | Inaccurate distributions for large clusters near bimodal regime; Scales $\sim N^{5.5}$ |
| SSA | Exact; Scales $\sim N^2$ | Slower than Mean-field CME for $N \lesssim 90$ |

Note that this model takes into account passive demethylation due to DNA replication in a continuous manner. That is, passive demethylation is assumed to be included in the *St*^−^ reactions.

### 2.2. CpG collaboration as a field effect

The collaborative, neighbor-CpG “field” effect is assumed to decay exponentially with distance, with lengthscale parameter *λ* (Fig. 1B). For CpG *i*, the demethylating field effect is given by

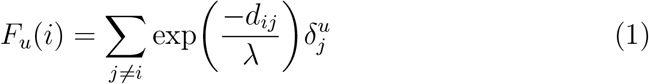

where the sum is over all other CpGs, *d_ij_* is the distance between the *i^th^* and *j^th^* CpG sites, *λ* is the interaction lengthscale, and δ*^u^_j_* = 1 if the *j^th^* site is unmethylated, and 0 otherwise. Similarly, the methylating field felt at the *i*th CpG is

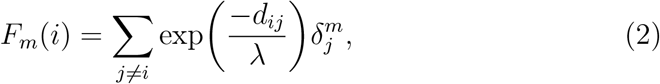

where δ*^m^_j_* = 1 if the *j^th^* site is fully methylated, and 0 otherwise. Thus, the combined rate (standard and collaborative) of *u* → *h* or *h* → *m* reactions at CpG *i* is *St*^+^ + *Co*^+^*F_m_*(*i*). Similarly, the *h* → *u* or *m* → *h* reactions at CpG *i* have combined rates *St*^−^ + *Co*^−^*F_u_*(*i*).

The complete stochastic propensities for each possible reaction to occur at an individual site *i* are given in Table 1.

### 2.3. Exact Chemical Master Equation Model

The chemical kinetic description of collaborative DNA methylation is essentially a network of first order reactions described by the Chemical Master Equation. Consider a network with *R* total reactions and let **x** denote the state of the system. The Chemical Master Equation is given by:

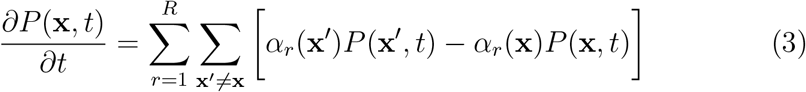

where *P* (**x***, t*) is the probability of being in a certain state at time *t*, **x**^′^ denotes a state different from **x** that is connected to it by a single-step reaction (that is, a change of methylation status of a single CpG), *α_r_*(**x**) is the propensity of reaction *r* occurring while in **x** [29]. The propensity corresponds to one of the *R* = 8 possible reactions, as in Table 1. For the DNA methylation model with *N* CpGs, let Ω = {*u, h, m*}*^N^* denote the complete state-space, where state **x** = (*x*_1_*, x*_2_*, …, x_N_*) ∈ Ω and *x_i_* ∈ {*u, h, m*} is the instantaneous methylation status of CpG *i*. We hereon refer to **x** as microstates.

In matrix notation, equation 3 can be expressed as

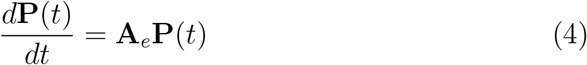

where **P**(*t*) is a vector containing the probabilities of being in each of the possible microstates in Ω at time *t*. **A***_e_* represents the rate matrix of the exact CME containing transition rates between each microstate. At steady state, equation 4 is

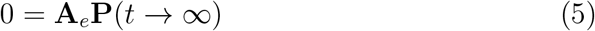

The non-trivial solutions **P**(*t* → ∞) to equation 5 give the probability distribution at steady-state, denoted **P***_S_*. The nonzero eigenvector corresponding to the nullspace of equation 5 is precisely this solution.

For a system with a single CpG site, there are only three possible microstates, and the reaction propensities are given solely by *St*^+^ and *St*^−^, as there are no collaborative effects. **P***_S_* gives the steady-state probability for the site to be in (*u, h, m*), with the single-site rate matrix denoted by **A***_e,_*_1_ and given by:

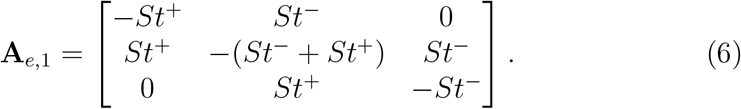

For *N* CpG sites, there are 3*^N^* microstates. Since the exact CME rate matrix **A***_e_* explicitly connects all of these microstates, **A***_e_* is a 3*^N^* × 3*^N^* matrix (though many elements are 0, since most microstates are not directly connected by a single reaction), and **P***_S_* is a 3*^N^* × 1 vector containing all combinations. The sums in Eqn. 3 are over all possible reactions in all possible microstates. By way of example, consider two particular microstates of the *N* = 6 system: Denote **x***_p_* = (*u, u, u, u, u, m*) as microstate *p* and denote **x***_q_* = (*h, u, u, u, u, m*) as microstate *q*. These two microstates are directly connected, since a *u* → *h* reaction at the first CpG transitions the system from *p* to *q*, and conversely a *h* → *u* reaction at the first CpG transitions from *q* to *p*. The off-diagonal matrix elements connecting these two states are given by

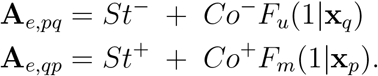

(Here, *F_u_*(1|**x***_q_*) signifies that the collaborative demethylating field “felt” at CpG *i* = 1 depends on the current status of all other CpGs in the system, i.e., it depends on **x***_q_*, and so on). Thus, the bottleneck here is that the fields *F_u_* and *F_m_* need to be separately computed for each of the 3*^N^* microstates, since they depend on the composition of the microstate and its relative positioning of *u* and *m* sites. Thus, exact solution of the CME requires enumeration of the full 3^2^*^N^* rate matrix.

### 2.4. Mean-field Chemical Master Equation Model

We developed [26] a simplified, mean-field approximation to the full CME model. In the approximation, microstates are grouped into macrostates based on the total number of CpGs of each type (*u, h, m*). For example, in this approach, (*u, u, u, u, u, m*) and (*u, u, m, u, u, u*) microstates of a 6-CpG-clusterm are lumped into the same macrostate, (5, 0, 1). Thus, **M** = (*N_u_, N_h_, N_m_*) represents a possible macrostate of the system, with the complete macrostate-space now given by Ω = *N_u_*∈ {0*, …, N* } × *N_h_*∈ {0*, …, N* − *N_u_*} (and then *N_m_* = *N* − *N_u_* − *N_h_*, as the number of CpGs is conserved). The mean-field CME can be constructed similar to above:

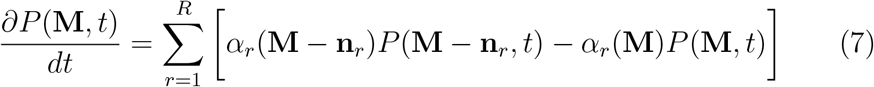

where *P* (**M***, t*) is the probability of being in a certain macrostate at time *t*, *α_r_*(**M**) is the propensity of reaction *r* at a given macrostate, and **n***_r_* is the stoichiometric change of the system from reaction *r*. For example, **n***_u_*_→_*_h_* = (−1, +1, 0) would describe the change in the number of (*u, h, m*) sites due to a methylation reaction *u* → *h*.

The propensities will be modified from equation 3 according to the mean-field approximation. Namely, the fields for each macrostate are computed as:

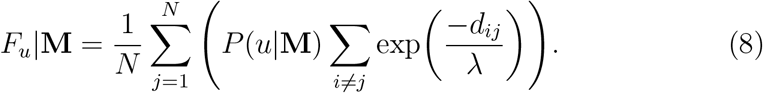

Here, *P* (*u*|**M**) is the probability for an individual site in the group of *N* CpGs to be in the unmethylated *u* state, given that the system is in macrostate **M**. Similarly,

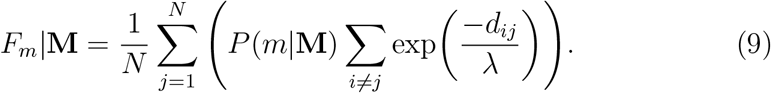

where *P* (*m*|**M**) is the probability for an individual site in the group of *N* CpGs to be in the fully methylated *m* state, given that the system is in macrostate **M**. Thus, Eqns. 8 and 9 average over all possible demethylating and methylating collaborative reactions, respectively, based on the collective averaged field exerted by CpGs on all others.

In vector-matrix form, equation 7 can be written as,

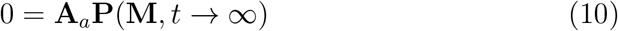

where **A***_a_* is the mean-field CME rate matrix. We denote the resultant steady-state probability vector over macrostates as *P_S_*(**M**). By way of example, consider two macrostates of the *N* = 6 system: **M***_y_* = (5, 0, 1) and **M***_z_* = (4, 1, 1), which are connected by a *u* → *h* reaction (and *h* → *u*, for the reverse transition). (Unlike the exact model, the mean-field model is agnostic to *where*, i.e. at which CpG site, this reaction occurs). The connecting matrix elements are given by:

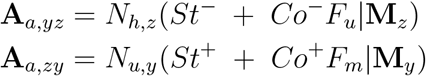

where *N_h,z_*and *N_u,y_* account for the multiplicity of ways that the relevant reaction can occur, depending on the number of *h* and *u* sites in macrostates *z* and *y*, respectively.

For *N* CpG sites, the number of macrostates is given by the triangular number

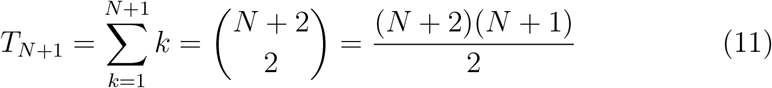

Thus the mean-field CME rate matrix **A***_a_* grows significantly more slowly with *N* than the exact CME rate matrix **A***_e_*.

Matrix methods are used to solve the CME; we use Scipy function **eig**. Note however that when solving only for the steady-state distribution, it is unnecessary to compute all eigenvectors, as only that associated to eigenvalue 0 corresponds to the stationary distribution. More efficient methods include **nullspace** and sparse methods such as sparse eigensolver **eigs** and sparse linear solver **spsolve**. In our study, the latter two methods achieved as much as a 10-fold speedup over **eig** for systems of O(10^2^) CpGs. However, the full set of eigenvectors can provide useful additional information in spectral analyses of dynamics, e.g., the computing of first passage time densities[30]. Thus, in our efficiency comparisions we use the **eig** function for CME computation.

### 2.5. Stochastic Simulation Algorithm Model

We apply the standard SSA[22] to the collaborative DNA methylation scheme. SSA uses a random-walk process to numerically solve the differential-difference equation (the CME). The algorithm and its application to collaborative DNA methylation is described below:

1. Define an initial time *t*_0_ and a maximum time limit *t_max_*. Define the positions of the *N* CpG sites. Define the initial microstate of the system using a numeric code *u* = 0*, h* = 1, and *m* = 2 (e.g. a fully methylated initial state is an array 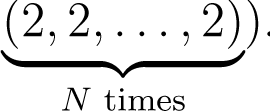. (If the simulation will be run to convergence at steady-state, then the initial condition is arbitrary).
2. Initialize the field effect matrix as exp(−D/λ), where D is the matrix with elements being the pairwise site-to-site distances dij.
3. Given the current system microstate, calculate the propensity matrix *α_ir_*, which stores the propensities of possible reactions *r* to occur at site *i*. Reaction index *r* runs over the eight types of reactions, for which associated propensities are shown in Table 1.
4. Sum over the propensities,

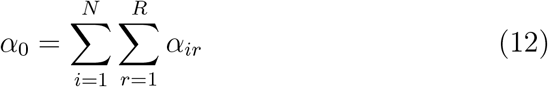

5. Generate a random number 0 *< r*_1_ ≤ 1. Compute the timestep *τ*

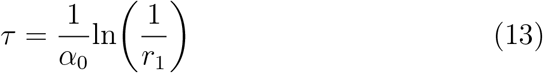

6. Generate another random number 0 *< r*_2_ ≤ 1. Find the smallest integer *k* such that

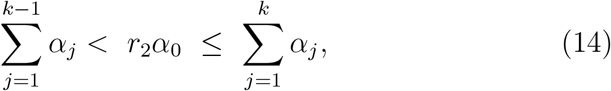

where *α_j_*is the “flattened” propensity vector holding *N* ×*R* propensities.

7. Convert the linear index *k* back to the row-column indices denoting the site *i* where the reaction *r* occurs. Update site *i* according to reaction *r* and update the time *t* by *t* = *t* + *τ*. Repeat the process from steps 3-7 until *t > t_max_* [29, 22].

### 2.6. Methylation distributions and summary statistics

Regardless of the computational method, methylation distributions are computed as the stationary probability over Methylation Ratio, or *P_S_*(*w*), where 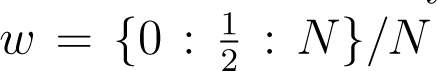 are the discrete, possible values of Methylation Ratio for a given system of *N* CpGs, since hemi-methylated sites count half as much as fully methylated sites:

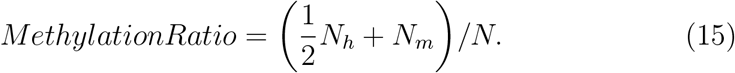

Thus, the Methylation Ratio is minimally 0 (for a group of CpGs all in *u* state) and maximally 1 (for a group of CpGs all in *m* state). This aids in comparing distributions across different system sizes. More than one system microstate (and, in the case of the mean-field model, macrostate) can have the same Methylation Ratio, thus the CME-computed complete state-space distributions are projected onto the Methylation Ratio bins for plotting. For the SSA approach, the methylation distribution is obtained by summing the relative time spent in each Methylation Ratio bin (mapped from individual microstates), harvested from a long trajectory after convergence to steady-state.

Another summary statistic to describe computed distributions is the bimodality metric. We devised a simple metric tailored to this system, given by:

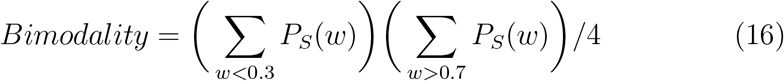

This metric uses cutoffs for low (0.3) and high (0.7) methylation, and its value ranges from 0 (for distributions with zero probability outside these cutoffs) to 1 (for distributions with all probability evenly polarized outside these cutoffs).

### 2.7. Simulation Convergence

To compute steady-state distributions from simulations, it is necessary to test for convergence. We used two types of convergence tests. For symmetrically parameterized models, (i.e., models with *St*^+^ = *St*^−^ and *Co*^+^ = *Co*^−^), an asymmetry-based convergence test was used. The simulation was run until a measure of distribution asymmetry fell below some specified tolerance level, *tol*. The asymmetry measure was given by:

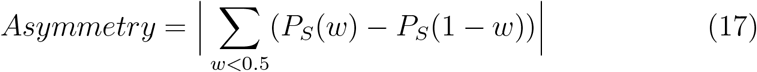

For asymmetrically parameterized models, the running *L*_1_-norm error was used to test convergence. The distribution computed cumulatively over time *t* = *κτ*, where *τ* here is a specified trajectory timelength, and *κ* is an integer, is compared to the previous increment by:

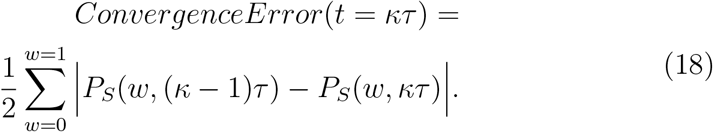

When this metric falls below a specified tolerance, it indicates negligible change of the distribution with increased simulation time.

### 2.8. Measuring accuracy of the mean-field approximation

To compare steady-state distributions obtained from different computational methods, we apply the *L*^1^-norm-based distance (also known as total variation distance),

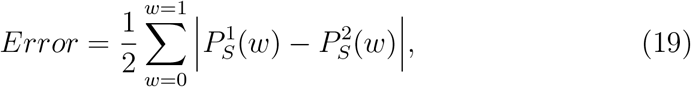

where here the *P_S_*(*w*) are obtained from two methods, denoted 1 and 2. The factor of 2 ensures that the Error metric is maximally 1 for two non-overlapping probability distributions. The choice of this metric allows practical comparison of error across different system sizes, which have different sets of allowed methylation ratios. In general, we use converged SSA simulations as benchmarks for the mean-field CME, since exact CME computation is impractical for all but the smallest system sizes.

### 2.9. Model parameters, time units, and data-driven assignment of standard rates

While we and others have estimated various methylation rates from varied data sources [13, 28, 21, 26], our focus in this manuscript was largely on numerical study of wide regions of parameter space, rather than on parameter fitting. Time units of model parameters herein are arbitrary, because steady-state distributions are sensitive only to the relative rates. Where we integrate the model with data, we assume *St*^+^ = 1, thus this parameter sets the intrinsic timescale of the system (i.e., all rate parameters are then in time units of (*St*^+^)^−1^).

We developed a data-driven assignment of the standard rate parameters as follows. The single-site standard model is analytically solvable via eqns. 5 and 6, giving *p_m_* = *N_m_/N* = (*St*^+^)^2^*/θ*, *p_u_* = *N_u_/N* = (*St*^−^)^2^*/θ*, and *p_h_* = *N_h_/N* = *St*^+^*St*^−^*/θ*, where *θ* = (*St*^+^)^2^ + (*St*^−^)^2^ + *St*^+^*St*^−^ (consistent with [19]). Ignoring collaboration, and using a data-derived estimate of the Methylation Ratio, *MR_data_*, and applying Eqn. 15, then *St*^−^ can be obtained by:

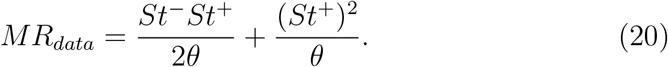

This equation connects the measured Methylation Ratio to the model parameters, assuming only the standard model. To this end, we use the Methylation Ratio as measured from only those CpGs that are assumed to be isolated; while no CpG is in reality perfectly isolated in the genome, we apply a threshold distance (O(10^2^ − 10^3^) bp) to identify isolated CpGs.

### 2.10. Data Availability

Software code for the three different methods along with data files related to mouse epigenome analyses are available at the following repository: https://github.com/Read-Lab-UCI/Computational-Methods-for-Stochastic-DNA-Methylation-Dynamics

## 3. Results

### 3.1. Different stochastic methods show collaboration-induced bimodality

We compared three computational approaches for modeling the stochastic, collaborative DNA methylation system: an exact CME, an approximate mean-field CME, and an SSA model (see Methods). Briefly, the exact CME computes the steady-state probability of all possible combinations of methylation states over all system sites, by enumerating the matrix of transition rates for all possible state-to-state transitions. For *N* CpGs, and neglecting symmetry considerations, there are 3*^N^* states (denoted microstates). Thus, enumeration of the rate matrix is impossible beyond ≈ 10 CpGs due to computer memory limits. The mean-field CME approximation averages over the group of *N* CpGs, essentially binning many microstates into fewer macrostates based on their overall methylation ratio (i.e., microstates with more *m* CpGs will tend to be lumped together, etc.). This is a type of mean-field approximation, since the field-effects driving collaborative reactions are now computed in an averaged way, rather than on the precise relative positions of *u* and *m* sites. The number of macrostates in the mean-field model increases as O(*N* ^2^), which enables reaching system sizes on the order of 100 CpGs (a relevant size for CpG islands: small CpG islands are a few hundred bp in length, while the average is around 1 kbp [31]). However, the mean-field model only computes probabilities over macrostates, rather than predicting behavior for individual CpGs.

In contrast, SSA simulation generates trajectories of the system behavior over time. It is an exact method, in the sense that the simulation is guaranteed to converge to the exact CME solution, if given sufficient time [22]. In contrast to the CME approach, the SSA requires low memory and does not require pre-enumeration of states or rates; the system traverses accessible microstates in a Markovian manner. Thus, SSA is often the method of choice for stochastic biochemical models[32], as it can handle large system sizes and/or high-dimensional state-spaces, but it can suffer from inefficiency.

We first validated the three different approaches on small methylation models. For the standard model, all three methods agree (Figure 1C). When collaboration is turned off, all sites behave independently, and the mean-field approximation is irrelevant. Thus, both CME approaches (exact and approximate/mean-field) produce the same methylation distribution, which agrees also with SSA. (In the absence of collaboration, the model also admits an analytical solution, which was also confirmed to match). With the model symmetrically parameterized such that methylating and demethylating rates are balanced, and no collaboration, the distribution is monomodal, centered at a methylation ratio of 0.5.

When collaboration is turned on, the distribution becomes bimodal, demonstrating how the positive feedback induced by CpG interactions leads to high or low methylation, i.e., collective behavior over the group of CpGs. In this parameter regime, SSA and Exact CME match well, but slight discrepancies appear in the distribution obtained from the mean-field CME model (Figure 1C).

### 3.2. Mean-field model runtimes scale more favorably with system size than exact models

A comparison of computational runtimes is presented in Figure 2. CME runtimes (both exact and approximate) are generally insensitive to the rate parameters of the model. This is because the bottlenecks for the CME approach are (i) the enumeration (by for loop) of all possible states and state-to-state transition rates, and (ii) the solution of Eqns. 4 (Exact) or 10 (Mean-field). Neither of these processes strongly depends on chosen rate parameters. Therefore, the CME runtimes depend directly on the number of states in the system. As expected, the least practical approach is the Exact CME: runtimes rapidly increase with *N* and were reasonably well-fit by the function (3*^N^*)^2^, which corresponds to a baseline estimation for scaling of solution of the 3*^N^* × 3*^N^* matrix. (All other methods were fit to a power-law (*N^k^*), but this model did not fit the Exact CME runtimes well).

**Figure 2:**
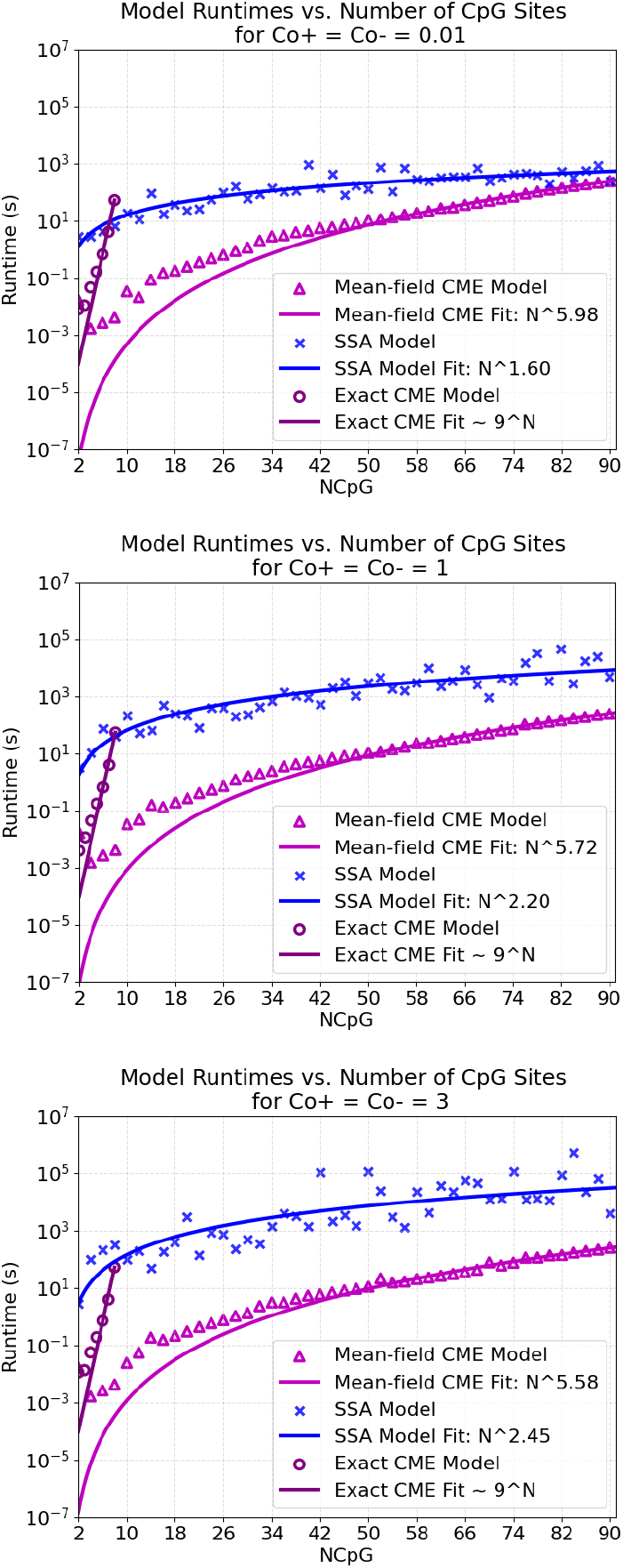
Exact CME, mean-field CME, and SSA model runtimes as a function of number of CpG sites. For all three models, standard reactions are set to *St*^+^ = *St*^−^ = 1. Lengthscale distance parameter given by *λ* = 30 with each CpG site equally spaced by 10 bp (i.e. *d* = 10 bp). **Top:** Model runtimes as a function of system size in an extremely low collaborative environment (*Co*^+^ = *Co*^−^ = 0.01). **Middle:** Model runtimes as a function of system size in a low collaborative environment (*Co*^+^ = *Co*^−^ = 1). **Bottom:** Model runtimes as a function of system size in a moderate collaborative environment (*Co*^+^ = *Co*^−^ = 3).

The Mean-field CME showed approximate power-law scaling with system size *N*, with a fitted exponent *k* of ≈ 5.58 − 5.98 across parameter sets. This scaling can be understood as follows. The number of macrostates *T* scales at least as quickly as O(*N* ^2^). More precisely, for a system of *N* CpGs, the macrostates are designated as comprising the collection of microstates whose methylation is characterized by {*N_u_, N_h_, N_m_*} (and *N_u_* + *N_h_*+ *N_m_* = *N*). Under this definition, the number of macrostates *T* is given by the triangular number, *T* = (*N* ^2^ + 3*N* + 2)*/*2 (see Methods). It follows that *T* ∼ *N^l^* where 2 *< l <* 3, and thus the baseline matrix computation scaling estimate would be *T* ^2^ = *N* ^2^*^l^* = *N^k^*. Thus 4 *< k <* 6. Note that, while the Mean-field CME method scales significantly more favorably than the Exact, it will also reach a memory limit, preventing treatment of systems with *N* ≳ 150.

In contrast to the CME methods, SSA runtimes depend on the simulation parameters as well as on system size. Runtimes for SSA generally increase with system size, because larger systems undergo more reactions per unit time. The variable-timestep SSA must explicitly track each of these reactions (although modified versions of the algorithm could help address this problem, e.g. [33, 34]). However, SSA simulations also converge at variable rates, depending on the parameters. Stronger collaborative rates leads to more extreme bimodality of the distribution, which is also associated with rare switching events between the two modes, of collective high methylation (mostly *m* across the *N* sites) and collective low methylation (mostly *u*). These rare events lead to long convergence times, due to the need to sample many such switching events in order to correctly estimate the relative probabilities in each mode. (Note that, in the symmetrically parameterized model, one knows *a priori* that the two peaks of the distribution should have equal probability. However, in general, the relative probabilities of the two modes is not known *a priori*).

We estimated runtimes for simulations of the symmetric methylation model in regimes of weak, moderate, and strong collaboration. These regimes are associated to no bimodality (and thus no rare events), moderate bimodality, and strong bimodality, respectively. In these regimes, the estimated power-law scaling exponent of SSA runtimes ranged from 1.60 to 2.45. These results demonstrate that, while SSA simulations for small system sizes are initially significantly slower than the Mean-field CME method, they scale more favorably with increasing system size, even for the strongly bimodal, rare-event regime.

### 3.3. Behavior of the model in different parameter regimes: Symmetric and asymmetric ways to achieve bimodality

We performed SSA simulations over a range of parameter regimes to comprehensively map the behavior of the methylation model. We explored the impact of varying both standard and collaborative rates on bimodality and average methylation ratio. For a model in which standard rates in the methylating and demethylating directions are balanced (*St*^+^ = *St*^−^), bimodality appears when collaborative rates *Co*^−^ and *Co*^+^ are nearly balanced, and sufficiently strong (Figure 3 Center).

**Figure 3:**
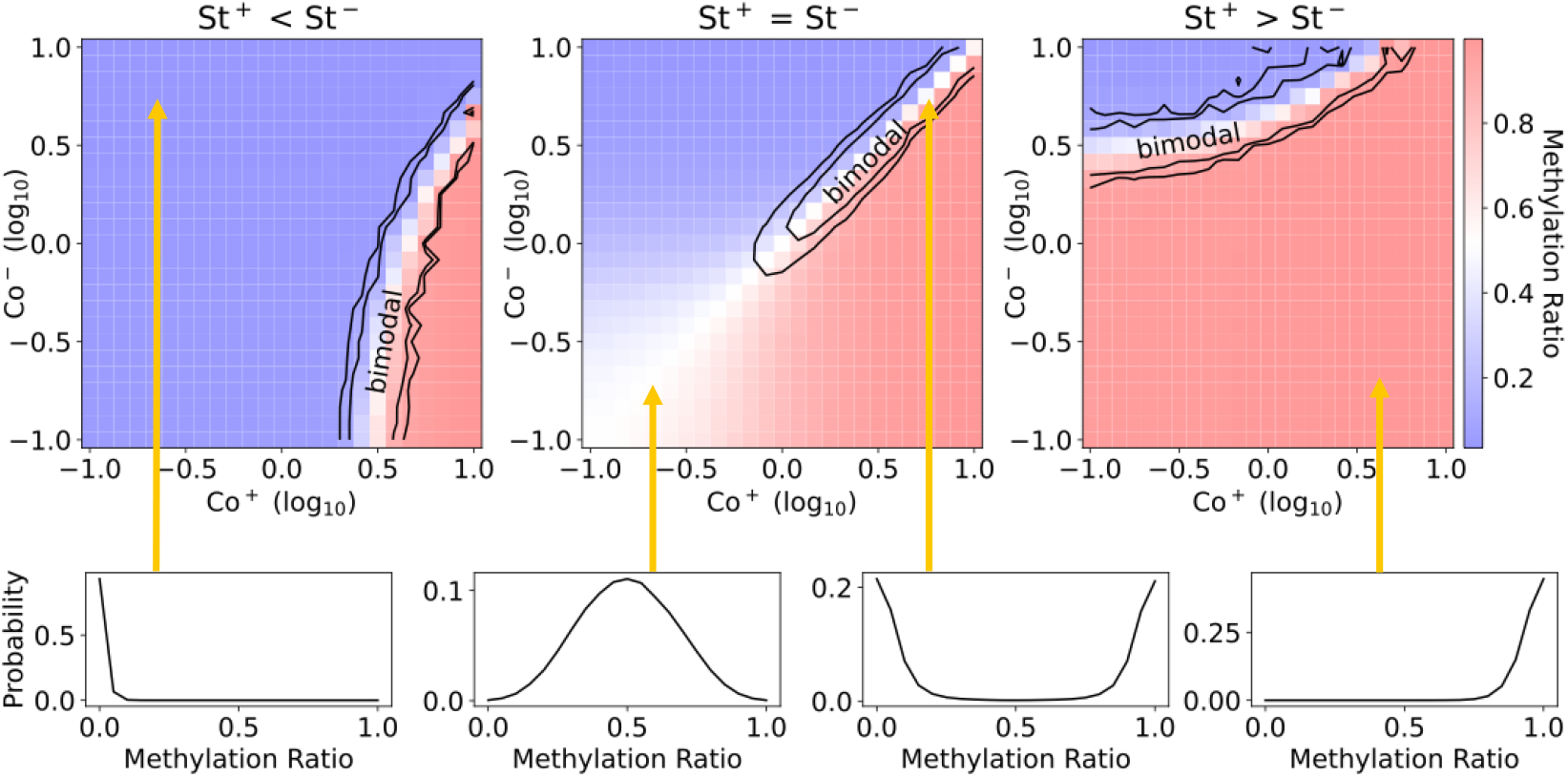
SSA simulation results: phase diagrams for the methylation dynamics model for a cluster of 10 CpGs (NCpG = 10, *λ* = 30 bp, d = 10 bp). Heat maps show methylation ratios (blue = unmethylated, red = methylated, white = intermediate) across the cluster as a function of collaborative reaction rates, *Co*^+^ and *Co*^−^, computed from the steady-state methylation distribution. Bimodality is present only within the contour lines, which correspond to values of [0.05, 0.15] for the bimodality metric. **Left:** Asymmetric model with standard reactions favoring demethylation (*St*^+^ = 0.1*, St*^−^ = 1.9). Bimodality is achieved in a region where collaborative reactions favor methylation. **Center:** Symmetric model (*St*^+^ = 1*, St*^−^ = 1). Bimodality is achieved for sufficiently strong and balanced collaborative reaction rates. **Right:** Asymmetric model with standard reactions favoring methylation (*St*^+^ = 1.9*, St*^−^ = 0.1). Bimodality is achieved in a region where collaborative reactions favor demethylation. **Bottow Row:** Representative converged distributions with arrows pointing to their corresponding points in parameter space.

Asymmetrically parameterized models also exhibit bimodality, when collaborative rates compensate for standard rates. For example, if *St*^+^ *< St*^−^, bimodality emerges in a region where *Co*^+^ *> Co*^−^. (see Figure 3 Left, and converse case, Right). Thus, positive feedback in the form of collaboration is only necessary in one direction (i.e., promoting only *m* or only *u*) for bimodality, so long as this positive feedback counters the flux in the other direction by standard reactions.

Overall, the necessary conditions for bimodality are sufficient collaboration in at least one direction, and net balance between the methylating and demethylating reactions.

### 3.4. Approximation error is greatest for large CpG clusters near mono-/bimodal transition

We investigated the effect of the size of the CpG cluster (*N*) and of CpG density (varying the inter-CpG distance (*d*)) on the methylation distribution, using both SSA and mean-field CME methods. In the asymmetric case, where we chose standard/collaborative rates favoring methylation/demethylation, respectively, both SSA and mean-field CME methods exhibit the same qualitative trend: Smaller, sparser clusters are methylated, while larger, denser clusters are unmethylated (Fig. 4, top). Bimodality emerges for certain size/density combinations in between these extremes. These results demonstrate how the system can use CpG sequence architecture to tune the bimodality and asymmetry of the methylation distribution in different genomic loci. The qualtiative trend matches data indicating that “less-interactive” CpGs (i.e., those that are more isolated, in CpG-sparse regions) tend to be methylated, whereas “interactive” CpGs (i.e., those in CpG islands, or more generally in denser/longer clusters) tend to be unmethylated [25, 26].

**Figure 4:**
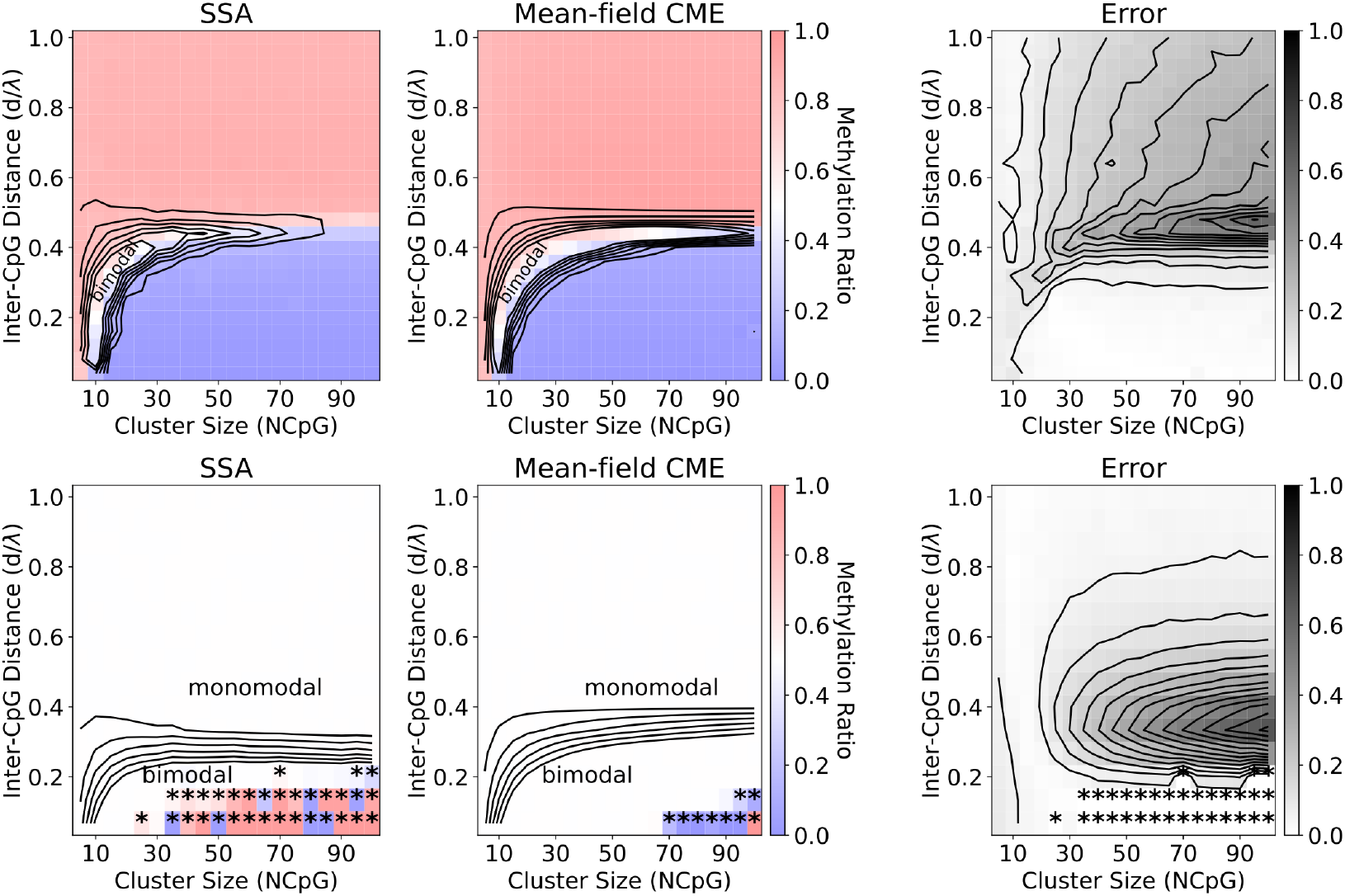
Comparison of phase diagrams over CpG cluster size and cluster density for the asymmetric model (top row) and symmetric model (bottom row), computed with either SSA simulation (left) or the mean-field CME model (center). **Top:** Model parameters are set such that standard reactions favor methylation, while collaborative reactions favor demethylation; values are fixed at *St*^+^ = 1*, St*^−^ = 0.35*, Co*^+^ = 1.2*, Co*^−^ = 2. **Bottom:** Model parameters are evenly balanced; values are fixed at *St*^+^ = 1*, St*^−^ = 1*, Co*^+^ = 1*, Co*^−^ = 1 (same parameters as Figure 2). Asterisks indicate lack of simulation convergence due to rare switching events (SSA) or unreliable numerical results due to ill-conditioned eigenproblem (CME). (Converged distributions have methylation ratio=0.5 (white) for the symmetric model.) **Both rows:** Bimodal region is shown by contour lines (0.1 to 0.6 in increments of 0.1 for levels of the bimodality metric). **Right panels:** For each cluster size/density, error is calculated between the SSA and Mean-field CME steady-state distributions by the L1-norm total variation distance. Error contour lines are at increments of 0.05.

In contrast to the asymmetric model, the symmetric model does not show a change of average methylation ratio with system size/density. Rather, different sizes/densities are always associated with symmetric methylation distributions, though characterized by differing extents of bimodality (Fig. 4, bottom). Bimodality is most pronounced for large, dense clusters. For both asymmetric and symmetric models, pronounced bimodality in the steady-state distribution corresponds to rare switching events in the SSA simulation trajectories.

We use the converged SSA simulations as a benchmark for validating the Mean-field CME method, using a normalized error metric (see Methods) (Fig. 4, right). Although the qualitative behavior over the parameter space from both methods is similar, quantitative errors in the computed distributions from the mean-field model can be high, particularly for larger CpG clusters near the bimodal-to-monomodal transition region.

For the symmetric model, a qualitative difference between the two methods is seen in the relative sizes of monomodal versus bimodal regions. The mean-field CME method predicts a larger bimodal region (bimodality metric *>* 0.6), extending, for example, past a density *d/λ* = 0.3 for clusters exceeding ∼ 60 CpGs. In this same region of parameter space from SSA, bimodality is weak (∼ 0.1 − 0.2, bimodality metric). This discrepancy appears to be due to the fact that the mean-field CME enforces spatial averaging across the whole cluster, whereas in SSA, local correlations can lead to patchiness–i.e., bimodality at smaller size scales that does not persist across the length of the entire cluster.

We noted numerical challenges for both methods in the extreme-rare-event (extremely bimodal) regime (denoted as asterisks in Fig. 4, bottom).

In this region, SSA convergence became prohibitively expensive. However, in this regime, the mean-field CME approach was also susceptible to numerical error. Since the system exhibits metastability, there is a near-degenerate pair of eigenvalues close to zero, making the eigenvalue problem numerically ill-conditioned. For either method, these issues resulted in the symmetric system (erroneously) showing methylation ratios different from 0.5.

### 3.5. Comparison to real-world methylomes

As the model provides a means of understanding how enzymatic processes control methylation as a function of both size and density of CpG clusters, we compared it to Whole Genome Bisulfite Sequencing data. We analyzed data from the Encode mouse reference epigenome (https://www.encodeproject.org/) [35]. We first analyzed the mouse genome (mm10) to identify clusters of CpGs. Following [25], clusters are defined using two parameters and two strict conditions: a cluster is a group of CpGs for which all nearest neighbor inter-CpG distances within the cluster are ≤ *d_max_*basepairs, and the boundary CpGs on the cluster edge must be ≥ *D_min_* basepairs from other (non-cluster) CpGs.

We show data-derived phase diagrams using a few representative clustering parameters (Fig. 5). The results show that large clusters exhibit characteristic densities. Small CpG clusters (*<* 15 CpGs) exhibit a broad range of densities, with inter-CpG distances ranging from 2 - >50 bp. In contrast, large clusters (extending from 15 to *>* 150 CpGs) tend to exhibit average inter-CpG distances of approximately 10 bp (although the precise mean distance depends on the parameters chosen in the cluster definition).

**Figure 5:**
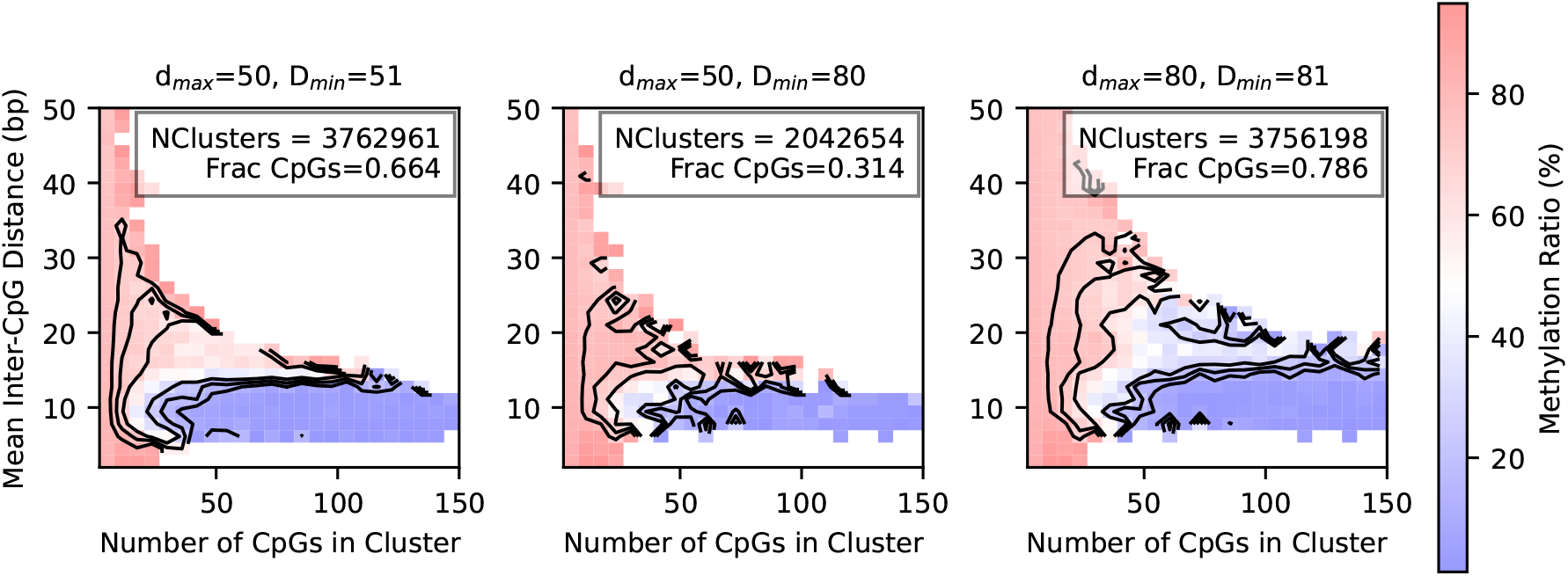
Data-derived phase diagrams from Whole Genome bisulfite sequencing of mouse large intestine (Encode accession ENCFF235GAI), using three different genomic CpG-clustering assignments (intra-cluster maximum CpG spacing, *d_max_*, and inter-cluster minimum spacing, *D_min_*). The center panel shows an assignment that enforces a larger inter-cluster gap, and thus better independence of individual clusters, but at the cost of losing a larger fraction of CpGs in the assignment. Inset text in each panel shows the total number of clusters found across all chromosomes, as well as the fraction of genomic CpGs that were assigned and thus included in the phase diagram. Bimodal region is shown by contour lines at 0.3, 0.5, 0.7 for the bimodality metric. Outer white regions indicate a lack of data; gridpoints are included in the heatmap where at least three clusters of corresponding *d*, *NCpG* are found.

The data show that dense, larger clusters are predominantly unmethylated, whereas the small clusters of variable density tend to be highly methylated (red versus blue regions in Fig. 5). In between these two regions, clusters exhibit bimodal methylation. This behavior is broadly in line with the asymmetric model, indicating that collaborative demethylation is stronger than collaborative methylation, and thus drives demethylation in large clusters. Fig. 5 shows a representative cell type from Encode, the mouse large intenstine (accession ENCFF235GAI). However, we computed qualitatively similar methylation phase diagrams across a range of mouse and human cell types.

Selection of parameters *d_max_*and *D_min_*for cluster assignment is not straightforward. Choosing a larger size of the flanking gap, *D_min_*, relative to the intra-cluster spacing *d_max_*, ensures that the cluster can be assumed to behave more independently of other CpGs, in line with the model (indeed, the model simulates a single, completely independent cluster, akin to flanking gaps of *D* = ∞). However, the downside of a too-strict condition on the relative sizes of *D_min_* and *d_max_* is that it can exclude the majority of genomic CpGs, since any instance of an intermediate inter-CpG-distance *d* (i.e., where *d_max_ < d < D_min_*) disqualifies all of the surrounding CpGs from being assigned to a cluster, regardless of their density. In the most permissive definition, we assign *D_min_* = *d_max_* + 1 bp, which ensures that the only CpGs neglected in the assignments are the individual “isolated” CpGs that are at least *D_min_* basepairs away from their nearest neighbors on both sides. This “permissive” definition captures many more CpGs but weakens the assumption of cluster independence. However, we found that the qualitative features of the phase diagrams were relatively insensitive to the cluster assignment parameters.

### 3.6. Complex phase diagram shapes

While the data-derived phase diagrams are qualitatively consistent with the asymmetric model of Fig. 4 (top), they also exhibit complex shapes that were not captured – namely, a “C-shaped” bimodal region. While the precise shape of the bimodal region is difficult to ascertain due to the lack of natural “samples” of clusters across the complete size/density space, there appears to be a localized region near cluster sizes below 20-30 CpGs, and densities of *d* ≈ 10 bp and closer, where increased density leads to higher methylation, in opposition to the broader trend (bottom left corner of panels, Fig. 5). We sought to investigate this feature using the model.

We first applied a data-driven estimation of the standard rate parameters, finding that *St*^−^*/St*^+^ = 0.36 for the mouse large intestine. (See Eqn. 20; for this estimate, we assumed that all CpGs more than 200 bp away from any neighbors are completely isolated, and thus well described by the standard model. The average methylation ratio across these CpGs in the dataset was found to be 74.66%). Next, we relaxed the assumption that both the methylating and demethylating collaborative lengthscales must be identical; instead we apply two different lengthscales *λ*^+^ and *λ*^−^, respectively. This then leads to a range of model behaviors as a function of the balance of collaborative strengths versus lengthscales, i.e., via *Co*^+^ and *Co*^−^ and *λ*^+^ and *λ*^−^ (Fig. 6).

**Figure 6:**
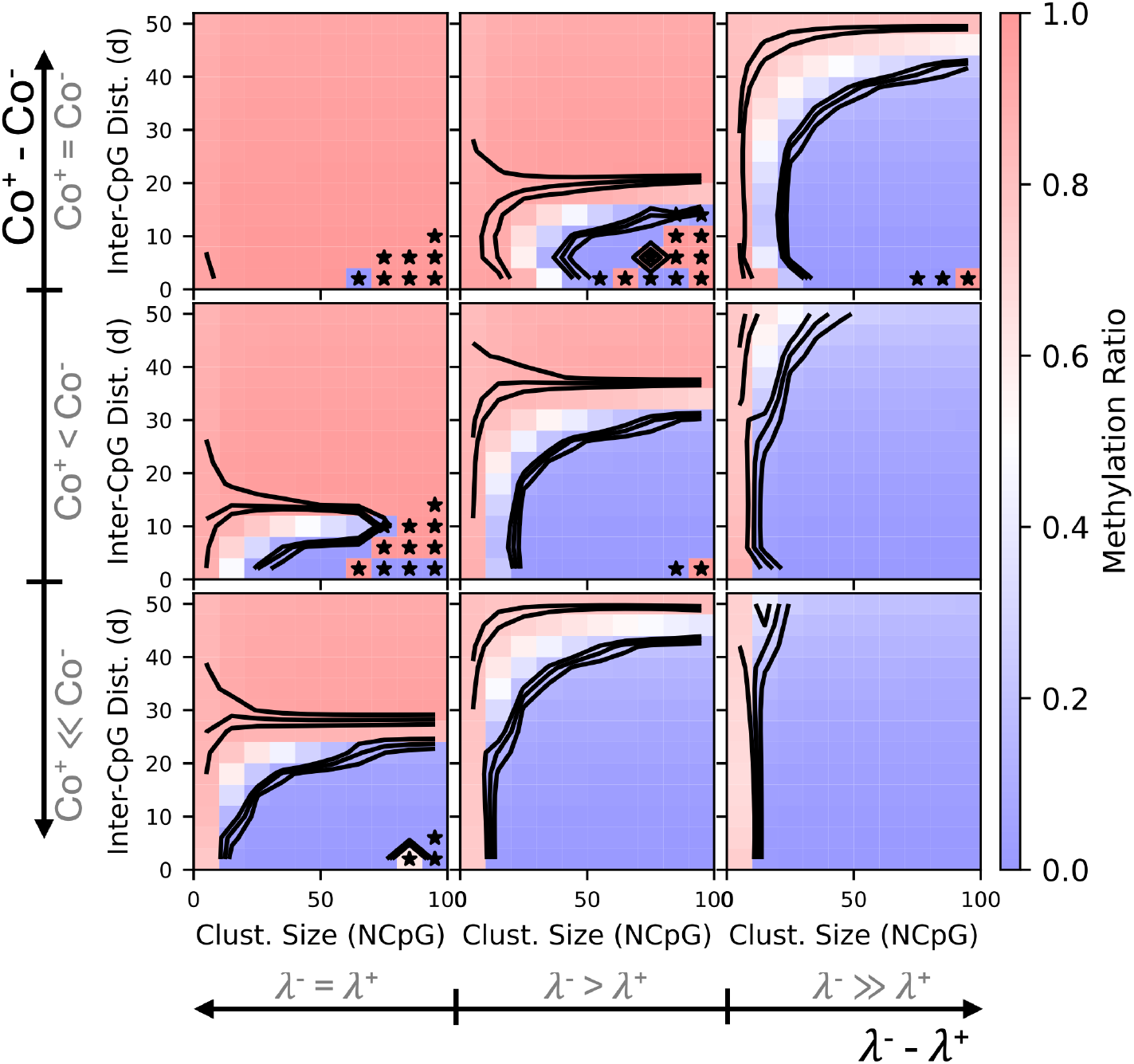
Comparison of phase diagrams as a function of CpG cluster size and density for nine different combinations of collaborative methylation rates and collaborative lengthscales, computed by the mean-field CME model. In all panels, *St*^+^ = 1, *St*^−^ = 0.36 (value estimated from data), *Co*^+^ + *Co*^−^ = 5.4 and *λ*^+^ + *λ*^−^ = 90 bp. **Rows:** Top to bottom, *Co*^−^ = 2.7, 2.97, 3.24. **Columns:** Left to right, *λ*^−^ = 45, 50.4, 55.8 bp. Bimodal region is shown by contour lines (0.1 to 0.3 in increments of 0.1 for levels of the bimodality metric). Asterisks indicate unreliable numerical results due to metastability (near-degenerate eigenvalues).

We found that choosing different lengthscales of collaborative methylation versus demethylation led to more complex features in the phase diagram. In particular, models with *Co*^+^ ≈ *Co*^−^ and *λ*^+^ *< λ*^−^ exhibit the “C-shaped” bimodal region, where small-to-intermediate sized clusters are unmethylated only at moderate densities. Here, clusters at low enough density (regardless of size) are methylated, because the effect of the standard reactions outweighs the collaborative reactions. However, clusters at very high density (*d* ⪅ 10 bp) are also methylated, because the strength of collaborative methylation is on par with demethylation at these lengthscales, i.e., as long as the total cluster extent is not too large. Only at larger cluster sizes does collaborative demethylation eventually win out.

All in all, these results demonstrate that real-world methylomes exhibit a complex dependence of cluster methylation on size/density of CpG clusters. The results also show that the model is capable of recapitulating these features, as long as the interaction lengthscales of the two opposing feedback loops are different.

## 4. Discussion

In this work, we comprehensively analyzed the behavior of a stochastic collaborative DNA methylation dynamics model, with a focus on how key features of methylation distributions—bi-versus mono-modality and overall levels of methylation (i.e., average methylation ratios)—are controlled by model parameters. Our results demonstrate how the interplay of standard (independent) and collaborative (interactive) enzyme-catalyzed “writing and erasing” of methyl marks, along with CpG locations and cluster sizes, determines these features of methylation distributions. A major challenge of such models is the *somewhat* collective behavior of model entities (here, CpGs): CpGs cannot be treated independently, but neither can they be treated as uniformly collective. Instead, the relative spatial positioning of CpGs, local interactions, and stochastic reaction events leads to rich dynamics that can also present computational challenges. We assessed the relative advantages and disadvantages, in terms of accuracy and efficiency, of different approaches (either simulation or Master Equation-based) to stochastic chemical kinetics modeling in application to the methylation dynamics model.

We also derived “phase diagrams”, analogous to those output from the nu-merics, from real-world methylation data. While the phase diagrams broadly show that larger/denser CpG clusters tend to be hypomethylated (a by-now well known phenomenon[36, 37, 38, 39]), our approach elucidates the relationship between methylation distributions and sequence-determined cluster architecture in a more fine-grained, quantitative way. We uncover a complex shape of the bimodal region, revealing a complicated (non-monotonic) trend of methylation with respect to CpG-density. The model reveals that this complexity is due to the competing feedback loops that promote (de-)methylation. Specifically, small CpG clusters show increased methylation with density, as long as standard rates favor methylation and collaborative rates for de-/methylation are similar. But if the interaction lengthscale of collaborative de-methylation is longer than that of collaborative methylation, then demethylation eventually wins, and larger clusters instead show decreased methylation with density.

Our mean-field model has the practical advantage of being more computationally efficient than stochastic simulations over the biologically-relevant size range of ⪅ 100 CpGs per cluster, especially when strong collaboration results in rare events. This enabled efficient study of the model behavior over a broad space of cluster and model parameters, thus enabling calculation of phase diagrams and easing integration with real-world data. Overall, we find that the mean-field approximation is generally accurate to the exact model in reproducing qualitative methylation patterns, but we reveal areas of the parameter space where approximation error is higher and hence care should be taken when the mean-field model is used to derive quantitative estimates. Interestingly, the real-world CpG clusters from the mouse genome seem to occupy areas of the cluster size/density space that are less susceptible to the mean-field approximation error. Since the mean-field model inherently assumes that the CpG cluster behaves as a unit (rather than allowing for spatial patterning within a cluster, as the full model does), this may suggest that evolutionary selection favors DNA-sequence-determined CpG clusters that are of the appropriate size/density, relative to enzyme-controlled interaction lengthscales, to function as a collective unit.

The model is a simplified version of the collaborative DNA methylation model as introduced in [14, 25]. We applied related mean-field modeling to aid in data analysis previously [26]. In contrast to those previous works, the current work more comprehensively explores the parameter space with additional focus on practical numerical considerations. We also here focus especially on the requirements for bimodality, which is thought to underlie the “epigenetic switch” function supported by DNA methylation. The mean-field model enabled rapid exploration of parameter space with generally high accuracy. A related mean-field model was developed by [19]; while their model differs from ours in various respects (e.g., focusing on neighbor-rather than distance-based interactions, and focusing on larger scale statistics, but not on bimodal patterning), they also found that a mean-field approach showed statistical accuracy to the full model, paving the way for better integration with data.

Our focus here was not on detailed parameter fitting of a particular dataset. However, our analysis of the phase diagrams of the mouse epigenome in various cell types points to a methylation system in which the route to bimodality is the “asymmetric” route with *St*^+^ *> St*^−^ and *Co*^−^ *> Co*^+^. This is reminiscent of our earlier findings from human cells[26], where we fit methylation parameters based on density-dependent methylation of individual CpGs, rather than on cluster phase diagrams. However, our findings here also point to a likely longer lengthscale of collaborative demethylation (*λ*^−^ *> λ*^+^), in which case the net collaborative demethylation effect can still be stronger in general (though not on short lengthscales), even when *Co*^−^ and *Co*^+^ are relatively balanced. In contrast, we and Lovkvist et al.[25] proposed that *λ*^+^ *> λ*^−^ in human cells; however, those prior studies employed a more complicated version of the model with many more rate parameters, including *h*−induced collaboration. The latter work[25] also proposed that human cells use the alternate route to bimodality (i.e., *St*^−^ *> St*^+^ and *Co*^+^ *> Co*^−^), whereas we propose that the high methylation of relatively isolated CpGs in both mouse and human cells suggests that *St*^+^ *> St*^−^. We anticipate that future, expanded applications of this type of modeling to mammalian genomes may shed further light on *in vivo* parameters.

A limitation of the model in its current form is the continuous replication approximation. Future studies could elucidate the impact on system dynamics of the mitotic cylce, in which replication is a periodic driving force of demethylation. Both the CME and SSA approaches are in principle amenable to including such a process into the model. Another limitation is the 1D-lattice model of CpG sites, which neglects the 3D structure of DNA. This limitation could also be addressed in future versions of the model, for example, by using spatial DNA-contact maps to inform the inter-CpG interactions. Such an extension could also be implemented within the current numerical frameworks, since it would affect only the function (here, simple exponential) that maps sequence-derived positions to interactions.

Although the SSA simulation approach was significantly more computationally expensive than the mean-field CME approach for clusters with *<* 100 CpGs, it also scaled much more slowly with *N* (O(*N* ^2.5^) for SSA as compared with O(*N* ^5.5^) for Mean-field CME). Thus, from an efficiency standpoint, the SSA model is only practical for more than 90 CpG sites, depending on the parameter regime. However, from an accuracy standpoint, care should be taken when applying the mean-field CME method, particularly for systems that are large relative to the interaction length-scale *λ*. Intuitively, the mean-field approximation breaks down for such large clusters, since the approximation allows CpGs to “feel” the effect of distant neighbors (in average), even when they should not according to the exponentially decaying field. We indeed observe this breakdown of the approximation for large system sizes near the bimodal regime.

Our analysis of computational efficiency is based on a traditional implementation of both SSA and CME methods. In the future, both approaches could potentially be sped up. For example, updated algorithms have been introduced to speed up SSA, such as tau-leaping and its variants [33, 34], which tackle the multi-temporal-scale nature of stochastic chemical kinetics simulations. Relatedly, the Weighted Ensemble approach has been demonstrated to speed up biochemical kinetic simulations of systems with rare events [40, 41]. As for the CME, the Finite State Projection (FSP) approach is applied to handle large (typically, infinite) state spaces by neglecting low probability (i.e., rarely visited) states [42], which could potentially address the memory-limit-issue that plagues the CME (even in mean-field). It is possible that an FSP-type approach could be applied to the DNA methylation model, although the model differs in important ways from typical biochemical kinetics models (spatial inhomogeneity of the system, rare events which could render rarely visited states critical to system dynamics, and the difficulty of knowing *a priori* which microstates are more probable). Alternatively, more sophisticated types of mean-field approximations could be adopted, for example, with averaging occurring over local neighborhoods rather than entire clusters. Symmetry arguments could potentially be used to reduce the size of the transition rate matrix (while bearing in mind that real CpG “lattices” are generally non-uniform). It may be possible to exploit rare events (i.e. timescale separation) to partition the rate matrix and thus reach larger system sizes with the CME [43]. Increased computational efficiency is desired, toward the goal of parameterizing models based on sequencing data across different cell types, or disease or therapeutic conditions.

## 5. Author Contributions

Both authors have made substantial contributions to the design and execution of the work. Conceptualization: ELR; Methodology: MAGL and ELR; Coding and investigation: MAGL and ELR; Data analysis: ELR; Writing, review, and editing: MAGL and ELR.

